# Challenges in solving structures from radiation-damaged tomograms of protein nanocrystals assessed by simulation

**DOI:** 10.1101/2020.09.18.298562

**Authors:** Ariana Peck, Qing Yao, Aaron S. Brewster, Petrus H. Zwart, John M. Heumann, Nicholas K. Sauter, Grant J. Jensen

## Abstract

Structure determination methods are needed to resolve the atomic details that underlie protein function. X-ray crystallography has provided most of our knowledge of protein structure but is constrained by the need for large, well-ordered crystals and the loss of phase information. The rapidly developing methods of serial femtosecond crystallography, micro-electron diffraction, and single-particle reconstruction circumvent the first of these limitations by enabling data collection from nanocrystals or purified proteins. However, the first two methods also suffer from the phase problem, while many proteins fall below the molecular weight threshold required by single-particle reconstruction. Cryo-electron tomography of protein nanocrystals has the potential to overcome these obstacles of mainstream structure determination methods. Here we present a data processing scheme that combines routines from X-ray crystallography and new algorithms we developed to solve structures from tomograms of nanocrystals. This pipeline handles image processing challenges specific to tomographic sampling of periodic specimens and is validated using simulated crystals. We also assess the tolerance of this workflow to the effects of radiation damage. Our simulations indicate a trade-off between a wider tilt-range to facilitate merging data from multiple tomograms and a smaller tilt increment to improve phase accuracy. Since phase errors but not merging errors can be overcome with additional datasets, these results recommend distributing the dose over a wide angular range rather than using a finer sampling interval to solve the protein structure.

## I. Introduction

Protein structure determination critically advances our understanding of biochemical mechanisms and the molecular basis of disease. X-ray crystallography has been the principal method of structure determination for decades, accounting for 90% of atomic models deposited in the Protein Data Bank [1]. However, two limitations constrain this method’s broader applicability. First, some proteins do not readily form sufficiently large and well-ordered crystals for characterization at synchrotron sources [2, 3]. Second, the phase information needed to solve a protein’s structure cannot be experimentally measured. Estimating this lost information requires experimental perturbations that some crystals are not amenable to, very high resolution diffraction data, or the availability of a homologous structure [4]. Such prior information is often limited for proteins that are difficult to crystallize, compounding the challenge of structurally characterizing new protein families.

The need for large crystals has been overcome in part by the development of techniques suitable for sub-micron sized nanocrystals. One of these methods is serial femtosecond crystallography (SFX), which relies on femtosecond-length X-ray pulses that are orders of magnitude brighter than synchrotron radiation [5]. SFX has significantly advanced structural studies of difficult-to-crystallize membrane proteins [6, 7] and crystals sensitive to radiation damage [8, 9] but suffers from low throughput. A second method is micro-electron diffraction (microED). This modality of cryo-electron microscopy (cryo-EM) takes advantage of the strong interaction between electrons and matter to enable data collection from crystals less than 1 *μ*m thick [10]. Similar to SFX, microED has enabled structural studies of amyloid proteins that do not readily form large, well-ordered crystals [11, 12]. However, neither method can measure the crystallographic phases, so this information must be inferred indirectly or supplied by other techniques [4].

Single particle reconstruction (SPR) is another cryo-EM modality that is increasingly effective for high-resolution structure determination. In SPR, projection images of purified proteins are recorded, aligned, and merged to determine the protein’s structure [13]. This method thus overcomes the main limitations of X-ray crystallography, bypassing the need for crystallization and retaining the phase information. Further, recent technological advances have enabled SPR to achieve reconstructions with high-resolution limits comparable to X-ray crystallography [13]. However, without the signal amplification that results from coherent scattering by a crystal, molecular weight is a limiting factor. As the object’s scattering mass decreases, errors in aligning projection images increase and attenuate the high-resolution signal [14, 15]. To date, the smallest macro-molecule solved by SPR to better than 4 Å resolution is a 40 kDa riboswitch, and only seven proteins <100 kDa have been solved to similar resolution [16]. By contrast, the median mass of proteins in the human proteome is 41 kDa [17], rendering SPR unsuitable for many proteins.

Cryo-electron tomography (cryo-ET) is a third modality of cryo-EM, and its application to nanocrystals could overcome the principal obstacles of mainstream structure determination methods [18]. In cryo-ET, a tilt-series of projection images is collected from a sample embedded in vitreous ice and reconstructed into a tomogram, a volume that contains 3D structural information about the specimen. This method has traditionally been used to study cellular ultrastructure, and the reconstruction approach of subtomogram averaging has recently determined near-atomic resolution (<5 Å) structures of select purified proteins and ribosomes *in situ* [19–21]. Like microED, cryo-ET is suitable for protein nanocrystals, with an expected optimal sample thickness of 50-300 nm [22, 23]. Like SPR, imaging rather than diffraction data are collected, so the phase information is retained during the experiment [24]. Further, collecting data in imaging mode provides a unique opportunity to spatially characterize and computationally correct for disorder [25, 26]. In 2D electron crystallography, this was achieved by a procedure called lattice “unbending” [26, 27]. Alternatively, disorder could be corrected in real space with subtomogram averaging algorithms [20, 28–32]. Another benefit of applying cryo-ET to nanocrystals is the potential to develop a hybrid method that combines high-resolution diffraction intensities from microED and low-resolution phases from imaging for structure solution. Even phases to intermediate resolution (~7 Å) should suffice to resolve *α*-helices and provide a robust starting model for phase extension [33, 34]. Development of this hybrid method should be straightforward, as sample preparation and the microscope are shared between the two techniques.

While electron imaging of 3D crystals has revealed lattice structure at nanoscale lengths [25, 35 36], to date structure determination has not been achieved from tomograms of such samples. In our first attempts to solve structures from experimental tomograms of nanocrystals, we found that low completeness was particularly severe at high tilt-angles and prohibitive to merging datasets. These observations prompted us to analyze the challenges of merging incomplete datasets and explore ways to improve data acquisition from nanocrystals. Additional difficulties, such as accounting for multiple scattering and the contrast transfer function, are anticipated during data processing [26, 37]. However, we focused on factors that predicted the ability to merge multiple datasets, as merging is critical to overcome the missing wedge of information inherent to tomography and loss of completeness due to radiation damage. Here we describe a data processing scheme to solve structures from tomograms of nanocrystals that leverages software from X-ray crystallography and algorithms we developed to handle challenges unique to tomography data. We also assess this workflow’s tolerance to the effects of radiation damage using simulated crystals. Our results recommend a data collection strategy that maximizes the angular spread of the reflections recorded in each tomogram to increase the likelihood of successfully merging datasets.

## II. Design of a data processing pipeline

Structure determination from tomograms of nanocrystals could leverage algorithms from crystallography or, alternatively, rely on subtomogram averaging [38] (see Discussion). Here we focus on the former approach. Since retention of phase information is unique to imaging methods, we emphasize the steps required to recover crystallographic phases from the tomogram’s Fourier transform. Our data processing pipeline leverages functions available in the SPARX [39, 40], DIALS [41, 42], and *cctbx* [43] software packages in addition to providing new algorithms developed specifically for tomographic data of crystal specimens. The steps are presented schematically in Fig. S1A and described in the following sections.

To develop this workflow and explore different ways of processing these data, we simulated tomograms of protein nanocrystals. We selected lysozyme in *P*1 symmetry (PDB ID: 6D6G) and a peptide inhibitor in *P*2_1_2_1_2_1_ symmetry (PDB ID: 4BFH) as model crystal systems to test distinct space group symmetries and protein folds. For each crystal system, the protein coordinates from the Protein Data Bank were tiled in Chimera to generate 3D nanocrystals containing 10 unit cells along each dimension [44]. Density maps were simulated from the nanocrystal’s atomic coordinates to 2.5 Å resolution using electron scattering factors with the *cctbx* software package and rotated to randomly-sampled orientations [43]. These intact volumes were projected into tilt-series spanning either a ±60° tilt-range with 3° increments or a ±40° tilt-range with 2° increments between projection images. No optical aberrations from the microscope, including the contrast transfer function, were simulated. The resulting tilt-series were reconstructed into tomograms using IMOD [45]. The workflow used to generate these simulated datasets is shown in Fig. S1B.

### A. Eliminating phase splitting

Real crystals are characterized by imperfections that result in loss of exact periodicity. Imaging introduces further non-idealities, such as interpolation errors from discrete sampling and truncation of crystal edges due to a finite field of view. These deviations from perfect crystallinity in real space result in spatially-smeared intensities and phase splitting at Bragg peak positions in reciprocal space (Fig. S2). Specifically, the phase values of pixels immediately surrounding peak centers shift by 180° between adjacent octants (Figs. 1A, S1). The split phases result from the presence of a circular discontinuity at the point considered the origin by the discrete Fourier transform, causing the phase to alternate sign between frequency bins in Fourier space. There is no such discontinuity for signals that are exactly periodic in the window of the Fourier transform, and hence no phase splitting for ideal (albeit finite) crystals.

**FIG. 1.**
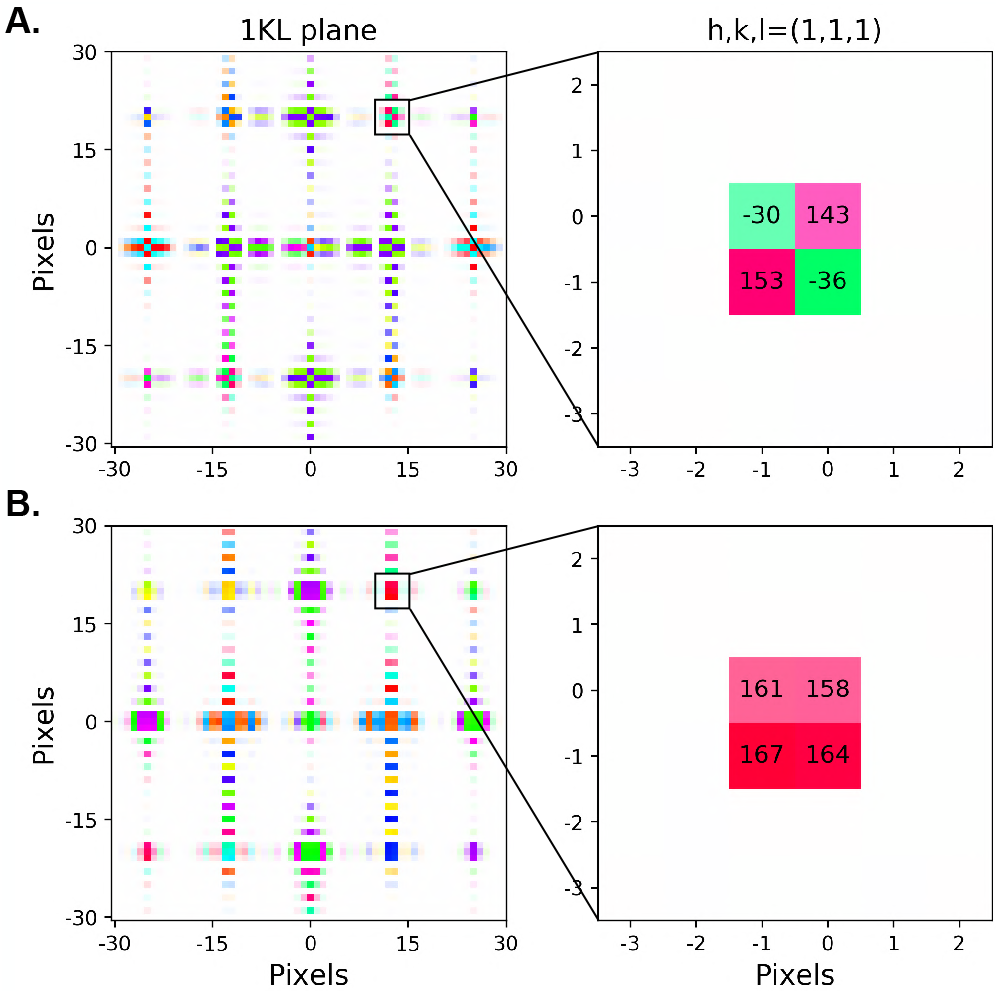
Elimination of phase splitting at Bragg peaks. (A) *Left:* The 1KL plane is visualized for a simulated crystal, which is imperfectly periodic because the unit cell dimensions do not span an integer number of pixels. The brightness and color of each pixel are determined by its intensity and phase values, respectively. *Right:* Inset of the (1,1,1) reflection. Pixels with intensity values within three-fold of the maximum intensity are visualized, and the phase values of these high-intensity pixels are noted in degrees. In (B), a tapering function was applied to the crystal. The density was centered by auto-convolution and shifted to the volume’s origin before computing the Fourier transform.

For imperfect crystals, we found that phase splitting could be eliminated by applying a symmetric tapering function in real space, followed by centering the density within the volume. Shifting the center of the resulting volume to the origin removes the circular discontinuity at the origin of the Fourier transform calculation. Here we chose a tapered cosine (Tukey) window with a tapering fraction of 0.5 and computed the translation needed to center the crystal density using the SPARX library’s auto-convolution function [39, 40]. Applying these pre-processing steps to crystals with imperfect periodicity eliminates phase splitting and results in a consistent phase value in the immediate vicinity of each Bragg peak (Figs. 1B, S2).

### B. Spot-finding and indexing

We adapted algorithms from the crystallography software package DIALS to index the Bragg peaks in the tomogram’s 3D Fourier transform [41]. Peaks were identified by scanning the Fourier transform along the x-axis (parallel to the axis of rotation), and high-intensity pixels in each slice were identified using the pixel thresholding algorithm described in Ref. [42]. Sets of contiguous bright pixels in adjacent slices were assembled into an initial spots list, which was then filtered according to the specified gain (the ratio of electrons per detector pixel to reported counts), resolution limits, and minimum and maximum number of included pixels [42].

Spot centroids were then mapped to reciprocal space. We used a 1d Fast Fourier transform algorithm to index the spots and estimate the crystal’s orientation and unit cell constants to within a magnification scale factor.[42, 46]. The provisional *P*1 cell constants were refined by enforcing constraints of the known space group symmetry. For experimental data, it is unclear whether the intensity information from imaging will be sufficiently accurate to determine the Laue class. Further, systematically forbidden reflections may be present due to multiple scattering or lie in an unsampled region of reciprocal space, preventing identification of screw axes [47]. However, symmetry could readily be determined by microED instead (see Discussion). Space group determination is critical for locating the crystallographic phase origin, as the phases do not reflect the space group symmetry until a crystallographic phase origin is found [47].

### C. Bragg peak fitting

Peak fitting involves assigning pixels to Bragg peaks, followed by integrating the intensities and averaging the phases of the assigned pixels. In X-ray crystallography and microED, profile fitting algorithms are used to integrate diffraction peaks [42, 48–50]. Profile fitting assumes a standard spot shape and models how each reflection is sampled based on its orientation with respect to the rotation axis, crystal mosaicity, and beam divergence [49]. In the case of tomographic data, however, a generic reflection profile cannot be assumed. Each reflection is only partially measured due to the large spacing between tilt increments and lack of continuous rotation (Fig. S3A), and at present we do not have adequate models to account for these effects. Developing a model to compensate for reflection partiality would improve the estimated intensities, but it would be challenging to devise a corresponding correction for the estimated phases.

Given these challenges, we implemented the following heuristic approach to determine reflection profiles from tomographic data. First, the pixel coordinates of each lattice point in the tomogram’s Fourier transform were estimated from the crystal’s indexing matrix. The reflection was discarded if it was predicted to lie in the missing wedge, or the volume outside of the ±60° or ±40° region spanned by the tilt-series. A spherical subvolume centered on each retained reflection within a specified radius was then considered; for the simulated crystals analyzed here, we chose a radius of 7 Å^−1^. Pixels were initially assigned to the peak if their intensities exceeded four standard deviations above the subvolume’s mean intensity (Fig. 2A-B). If multiple sets of noncontiguous pixels were found, the set with the centroid nearest to the predicted peak center was retained. The reflection was discarded if the observed peak centroid exceeded a distance of 2 reciprocal pixels to the predicted peak center. Such discrepancies between the ideal and observed Bragg positions resulted from peak centers being poorly sampled due to the large angular spacing between tilt increments. These partially-measured reflections were rejected because the observed phase values were frequently shifted from the expected values by 180° (Figs. S4-5).

**FIG. 2.**
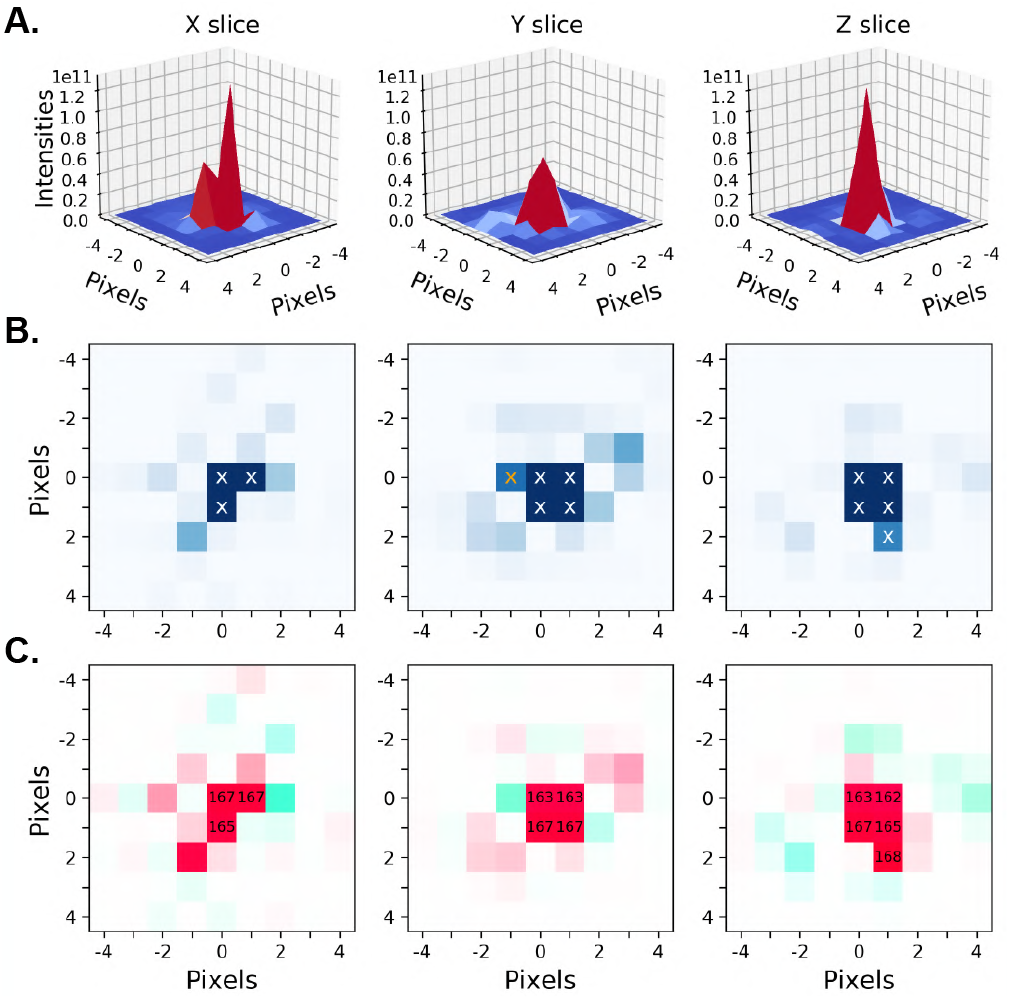
Peak fitting for a representative reflection. (A)The intensity profiles of slices are visualized along the indicated direction and centered on the (−2,3,−8) reflection of a simulated crystal tomogram. In (B), the intensities are shown in 2D, with high-intensity pixels assigned to the peak marked by an X. In (C), the phases in the vicinity of the peak are visualized using the same color scheme as in Fig. 1. The phase values of pixels retained as part of the peak are noted in degrees. The high-intensity pixel marked by a yellow X in (B) had a phase value of 17° and was discarded as an outlier.

To estimate the background intensity, the same subvolume used to assign peak pixels was considered. Both the pixels assigned to the peak and any contiguous pixels two standard deviations above the subvolume’s mean intensity were masked, with unmasked pixels considered background. This threshold was less conservative than the one used for assigning pixels to the peak to reduce the likelihood of including high-intensity outliers in the background region. We then estimated the values of the masked pixels by trilinear interpolation to approximate the background intensity beneath the peak. We used interpolation rather than assuming a constant background to better account for the anisotropic structure of the non-Bragg intensity, which results from the measured data being oriented along skew slices of finite thickness that are commonly separated by an angular increment of 2 or 3°. As a result, the orientation and magnitude of the background vary among peaks, depending on the peak’s position relative to the planes of sampled signal in Fourier space (Fig. S3). The estimated background values were then subtracted from the original intensities of the corresponding pixels.

The assignment of pixels to Bragg peaks was then refined using the phase values (Fig. 2C). Specifically, the intensity weighted standard deviation of the assigned pixels’ phases was computed. If this metric exceeded a specified threshold, outlier pixels were iteratively removed until this threshold was reached. Empirically we found that a threshold of 15° enabled the recovery of accurate phase values. Ensuring consistent phase values within the peak is critical, as the phase can change sharply outside the Bragg peak (Fig. 2C). Finally, the reflection’s intensity and phase were respectively estimated from the sum of the intensity values and the intensity-weighted mean of the phase values of the retained pixels.

Both the intensity and phase components of the Bragg peaks were fit using tomograms pre-processed as described in Section IIA, which eliminated phase splitting. Although application of a tapered cosine function in real space reduced the intensities, it also prevented sharp edges in the window of the Fourier transform from being convolved with the shape of the Bragg peaks in the frequency domain. As a result, the reflection profiles were more spherical and a higher number of Bragg peaks were retained by the peak-fitting algorithm described above, which benefited from this rounder shape.

### D. Merging datasets

In cryo-ET, the experimental geometry limits the accessible tilt range to ±70° relative to the sample’s untilted orientation [38, 51]. In principle, this rotation range should suffice to collect a complete dataset from a single crystal for most point groups [52]. In practice, however, 140° is an overestimate of the useful rotation range because intermediate to high-resolution information is lost both at high tilt angles due to increased specimen thickness and in images recorded late in the tilt-series due to accumulated dose [53]. Further, a popular data collection strategy in cryo-ET uses a 3° tilt increment for a tilt-range of ±60° [53], so a fraction of Bragg peaks in the nominal tilt-range will fall between recorded images and not be sampled. The size of this fraction is anticipated to vary between samples and depend on factors like mosaicity and the resolution range. Achieving high completeness thus requires merging data from multiple crystals in different orientations.

Merging phases from different crystals requires positioning the datasets on a common phase origin. Unless this is also a crystallographic origin, the phases of symmetry-equivalent reflections are not related and the phase data effectively have *P*1 symmetry. Here we established a common phase origin by treating one crystal as reference and shifting the phase origins of the remaining datasets to this reference origin (Fig. 3). The fractional unit cell row vector, **u**, that shifts the phases of a second crystal to the reference origin was determined by minimizing the intensity-weighted mean residual between the reference phases and the shifted phases of the second crystal:

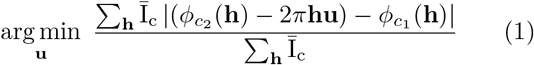

where the minimization is performed over all discrete values of **u** that fractionally sample the unit cell in real space according to a specified interval along each cell dimension. The sum is over the shared reflections between datasets *c*_1_ and *c*_2_, **h** is a column vector of Miller indices, and 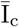 is the mean intensity for reflection **h**. Eq. 1 is similar to the phased translation function used in molecular replacement searches, though here we compute intensity-weighted phase residuals rather than calculating complex structure factors [54]. The term in parenthesis, 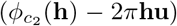, describes how the phases of *c*_2_ change when its phase origin is shifted by **u**. This expression was then used to position the phases of all reflections of *c*_2_ on the reference origin. To merge the next crystal, the reference phase set was updated to include all reflections recorded in the original *c*_1_ and *c*_2_ datasets. The reference phase set thus expanded with each merge while the reference origin remained roughly fixed, with only minor adjustments after averaging the phases from different crystals. Datasets were merged in the order the maximized the number of shared reflections between the reference data and the dataset being merged at each step. Once the final dataset was merged, the phase of each reflection was estimated as the intensity-weighted mean phase for all observations of that reflection.

**FIG. 3.**
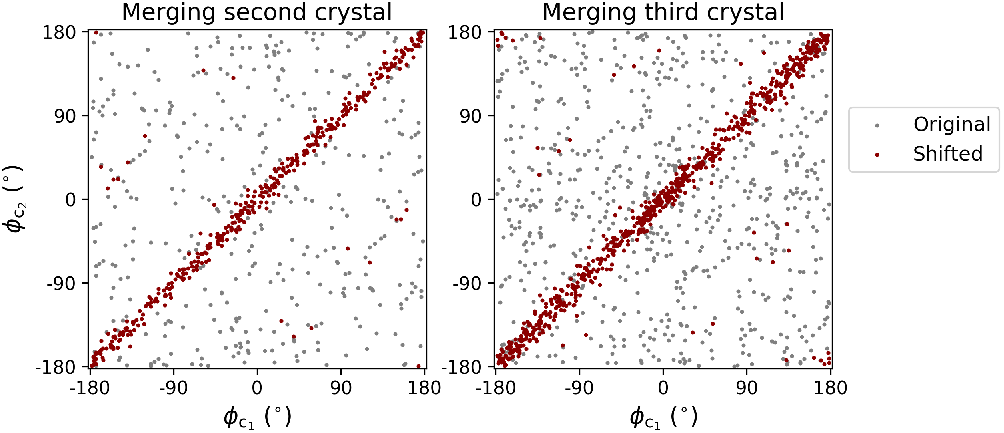
Identifying a common phase origin. The phases extracted from tomograms of three randomly oriented, simulated crystals were merged. Each panel shows the relationship between the phases of reflections shared between the reference 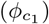 and added datasets 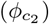 before (grey) and after (red) shifting the latter to the reference origin determined by the first crystal. With each additional dataset, the number of reference reflections increases, while the origin remains fixed.

Intensity information is unaffected by the choice of phase origin due to translational invariance. For consistency, however, we also treated the intensities as having *P*1 symmetry during merging. Intensities from a second dataset, 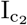, were uniformly scaled to the first dataset, 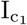, by minimizing the sum of squared residuals:

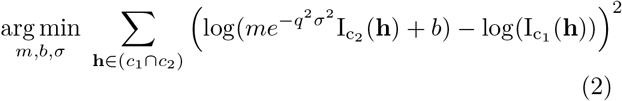

where *q* is the magnitude of the reciprocal space vector associated with reflection **h**, the exponential term is the Debye-Waller factor [55], and least-squares optimization is used to determine the scaling parameters *m*, *b*, and *σ*. We found that the use of logarithmic residuals improved the stability of the optimization algorithm. The next dataset was then scaled to the averaged intensities of the original *c*_1_ and *c*_2_ datasets. Once all datasets were merged, each reflection’s intensity was computed as the mean of all observations of that reflection.

### E. Locating a crystallographic phase origin

As noted above, it is unlikely that the phase origin of the merged dataset will coincide with a crystallographic origin consistent with the crystal’s space group. In 2D electron crystallography, the crystallographic origin was found by testing fractional cell positions in the plane of the repeating unit for fulfillment of phase constraints imposed by symmetry [47, 56]. If the plane group was not known in advance, this origin refinement procedure was performed for every possible plane group, and the plane group that yielded the lowest phase residual was selected. Crystal symmetry determination is easier for 2D crystals, however, as there are only 17 plane groups in contrast to 230 space groups [47]. Here we assumed that the space group was already known and extended the method of origin refinement to 3D crystals as follows.

The unit cell was sampled using a real space grid with equally-spaced nodes along each dimension, and each node or fractional cell position was considered a candidate origin. The phases of the merged data, *ϕ*_0_, were shifted by the fractional cell vector, **u**, to this origin:

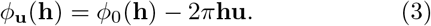

We then evaluated the following three metrics. The first metric corresponded to the phase residual of symmetry-equivalent reflections. The mean phase value for a given reflection **h** can be estimated from its symmetry-equivalent reflections as follows [57]:

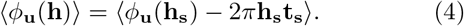

where the symmetry-equivalent reflections, **h** and **h**_**s**_, are related by the translation vector, **t**_**s**_, and the phase origin is **u**. We then computed the symmetry phase residual as the sum of the phase differences between this mean value and independent observations from the symmetry-equivalent reflections:

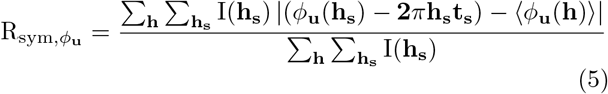

where the double sum enumerates the set of unique reflections and, for each, the set of symmetry-equivalent reflections. The residual was intensity-weighted to favor stronger reflections.

The second metric considered the difference between centric reflections and their expected phase values. Centric reflections satisfy the condition **hR** = −**h**, where **R** is the rotation matrix associated with reflection **h** [57]. When reflection data are positioned on a crystallographic phase origin, the phases of centric reflections are restricted to a limited set of possible values. We computed the intensity-weighted mean residual between the observed phase values at each candidate origin **u** and the expected phase values for these reflections as follows:

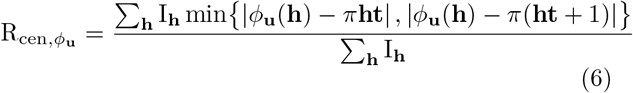

where **t** is the translation vector associated with reflection **h**, and the sum is restricted to the observed centric reflections [58]. This metric was omitted for datasets that did not contain any centric reflections.

For the third metric, an electron density map of the unit cell was computed from the intensities of the merged dataset and the shifted phases, *ϕ*_**u**_, using the *cctbx* library [43]. The skew of the density values was evaluated, as the density distribution of macromolecular crystals exhibits a positive skew in contrast to the Gaussian distribution characteristic of random maps. This distinction is used to judge map quality during automated structure solution after experimental phasing [59]. The advantage of this metric is that it uses all available reflections, rather than just the subset of either centric reflections or reflections with high multiplicity. For consistency with the phase residual metrics, the negative of the skew was computed such that lower values indicated more probable origins.

These three metrics were normalized and summed with equal weighting to score each candidate origin given by the fractional cell vector, **u**. The fractional cell position with the lowest combined score was selected, and the phases of the merged dataset were shifted to this origin. The intensities and shifted phases were then reduced to the asymmetric unit, and symmetry constraints were imposed. Specifically, the phases of symmetry-equivalent reflections were mapped to their expected values in the asymmetric unit (see Eq. 4) and the intensity-weighted mean phase value was computed. The intensities of symmetry-equivalent reflections were averaged.

As an alternative approach, individual datasets could be positioned on a crystallographic origin prior to merging. In this case, the merging procedure would require a search over a limited number of positions rather than the entire unit cell, as space group symmetry restricts the number of crystallographic origins. However, we found that the process of merging two datasets was slightly more robust than positioning individual datasets on a crystallographic origin. We compared each strategy using simulated datasets of a *P*2_1_2_1_2_1_ crystal with unit cell dimensions a,b,c=16.2, 29.1, 47,7 Å generated with a range of mean phase errors (0-40°), completeness (10-40% prior to accounting for space group symmetry), and relative B-factors (described in more detail in Section III; see Eq. 7). The searches for the common origin between datasets and crystallographic origin were both performed using a sampling interval of 0.2 Å; the phase origin was considered to be correctly found if the distance between the correct origin and the origin found by the algorithm was less than twice the sampling interval. We observed a 4% higher success rate for finding a common phase origin between two datasets than locating a crystallographic origin for each individual dataset (Fig. S6A). Further, identifying a crystallographic origin was successful in 8%more cases when the procedure was performed on the merged data from two tomograms than on a single tomogram (Fig. S6B). In contrast, the success rate for merging two and three datasets was similar (Fig. S6C). These trends suggest a modest advantage to merging datasets to a common phase origin prior to finding a crystallographic origin, as the latter procedure appeared to be more sensitive to the number of available reflections.

## III. Tests of robustness to radiation damage

We validated the full workflow presented in Section II (Fig. S1A) by evaluating the accuracy of the reflection data recovered from simulated tomograms. The processed data were compared to phases and intensities computed directly from the atomic model, in addition to reflection data recovered from “intact volumes” processed in the same manner as the simulated tomograms. Intact volumes refer to rotated crystal densities generated in the same manner as the simulated tomograms (see Section II), except without projection into tilt-series and reconstruction into tomograms. This comparison between intact volumes and simulated tomograms enabled assessing the inaccuracy and loss of completeness introduced by tomographic sampling beyond the baseline interpolation errors from simulating and rotating the crystal densities onto a discrete grid. For each structure (6D6G/4BFH) and type of volume (intact volume/tomogram), performance was judged based on merging five datasets. This merging procedure was repeated ten times with unique sets of five datasets.

Metrics evaluating the quality of the recovered reflection data are shown in Table 1. The information from the intact volumes was consistent with the reference phases (R_ref,*ϕ*_ <2.5°) and intensities (CC>0.97) computed from the initial atomic models. For the simulated tomograms, merging multiple datasets was required to achieve high completeness. When the tilt-range spanned 120°, or 67% of reciprocal space, the data recovered from each tomogram were only 40% complete in the absence of internal symmetry (Table 1). The unexpectedly low completeness stems from the large angular increment of the tilt-series: many Bragg peaks lie between tilt images and were either not observed or discarded as poorly sampled. Generating tomograms using a ±40° tilt-range with 2° increments rather than a ±60° tilt-range with 3° increments resulted in a slight decrease in the overall completeness (~5%) but a modest improvement in the accuracy of the reflection data due to the finer sampling (Table 1). Merging tomograms and the presence of internal symmetry also improved the accuracy of the recovered phases, which showed good agreement with reference (R_ref,*ϕ*_ of 6.2° and 2.8° for the *P*1 and *P*2_1_2_1_2_1_ crystal systems, respectively, when the tilt-range spanned ±60°).

**TABLE I.**
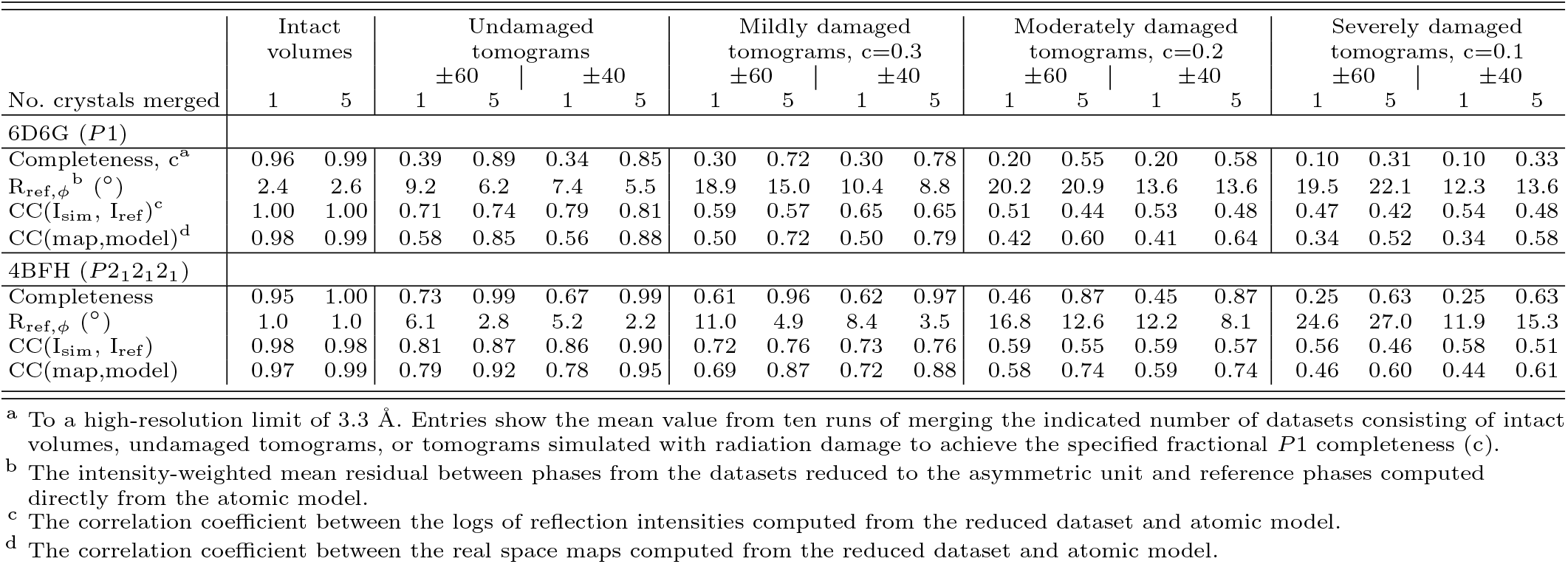
Quality of merged data

In real space, loss of information due to the missing wedge leads to elongation of density in the direction of the beam [60]. These anisotropic effects were apparent in density maps computed from single tomograms of the *P*1 crystal, with continuous density along parallel bands and gaps along the protein backbone between these streaks (Fig. 4A). Incorrect connectivity was less pronounced in density maps computed from single tomograms of the orthorhombic crystal, as the presence of internal symmetry mitigated the directionality of the missing wedge effect (Fig. 4B). In both cases, merging multiple crystals improved the correlation with the reference map (Table 1).

**FIG. 4.**
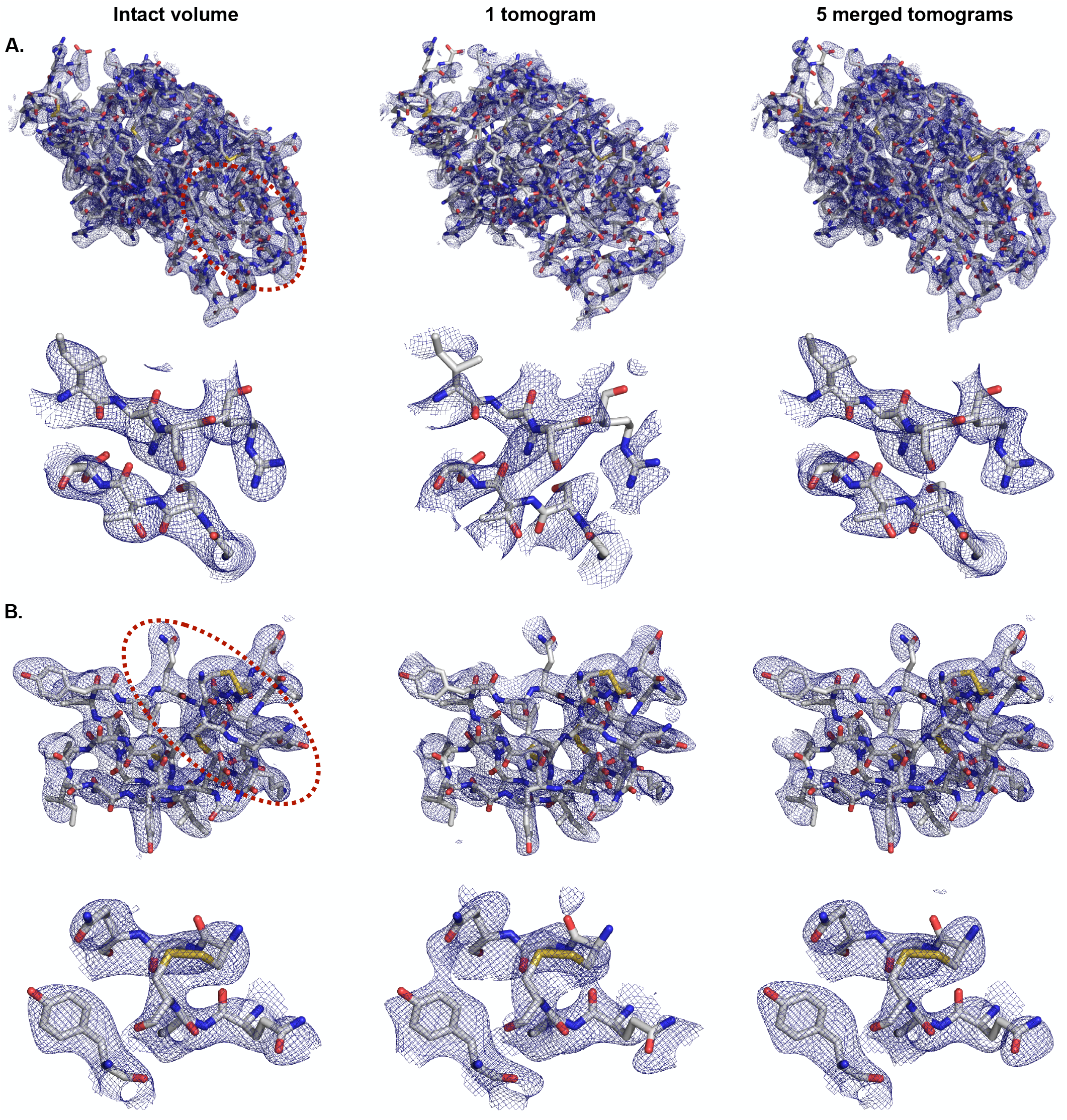
Merging undamaged tomograms recovers the reference density. Density maps were computed from reflection data for a representative intact volume (left), one tomogram (center), or the merge of five tomograms (right). The simulated crystal system had either (A) triclinic or (B) orthorhombic symmetry. Each panel visualizes the map for the entire protein (upper) and the subset of residues circled in red (lower).

Radiation damage is a limiting factor in cryo-ET and results in loss of information, particularly at high-resolution, as data collection progresses [61]. To assess how this impacts recovery of the reference structure, we next applied the workflow to tomograms of these crystal systems with simulated damage. Damage events were assumed to occur at random sites in the crystal volume, and each “hit” was modeled as a blurring of the local density [62]. The blurring was performed by replacing a cubic subvolume of length 5 Å around each selected site with its Gaussian-filtered copy, using a standard deviation of 1 Å along each axis for the 3D Gaussian kernel. Following a dose-symmetric tilt scheme [53], an equal number of hits was applied to the randomly-oriented crystal before computing each projection image to mimic a linear increase in absorbed dose across the tilt-series. Tomograms were reconstructed from these damaged tilt-series, and those that could not be indexed to the correct space group were discarded. For both crystal systems, we calibrated the total number of hits, or effective dose, that on average yielded tomograms with a completeness of 30, 20, or 10% before accounting for space group symmetry, compared to the ~40% *P*1 completeness observed for undamaged tomograms (Table 1). To focus on the tolerance of data processing to low completeness, ten unique sets of five tomograms were merged if the completeness of each tomogram and the set’s average was respectively within 2% and 1% of the target completeness. Metrics evaluating the average quality of the individual tomograms and merged datasets are reported in Table 1.

For both crystal systems, the reference structure was consistently recovered by merging tomograms that were each 30% complete prior to accounting for space group symmetry. Merging improved the accuracy of the phases and resolved connectivity errors observed in real space maps computed from a single damaged tomogram (Table 1, Fig. 5). While an initial completeness of 20% could also be tolerated by the orthorhombic system, results for the triclinic crystal were variable, with the phase accuracy frequently decreasing during the course of merging despite the increase in multiplicity (Fig. S7). For both crystal systems, an initial completeness of 10% per tomogram was prohibitive to recovering the reference structure. Despite the modest increase in cross-correlation with the reference map, merging reduced phase accuracy under this starting condition, indicating that a common phase origin was not correctly found (Fig. S7). The density of the merged tomograms showed the wrong connectivity, including smearing of density in the solvent region between neighboring chains and gaps along the backbone (Fig. 5, right). In several cases, the merging procedure visibly aligned the missing wedges of the tomograms being merged (Fig. S8), suggesting that the shape of the missing wedge rather than the reflection data dominated the signal. Similar trends in the relationship between initial completeness and failure to merge were observed when damaged tomograms were reconstructed from tilt-series spanning either a ±60° tilt-range with 3° increments or a ±40° tilt-range with 2° increments, despite the lower initial phase errors of the latter (Table 1).

**FIG. 5.**
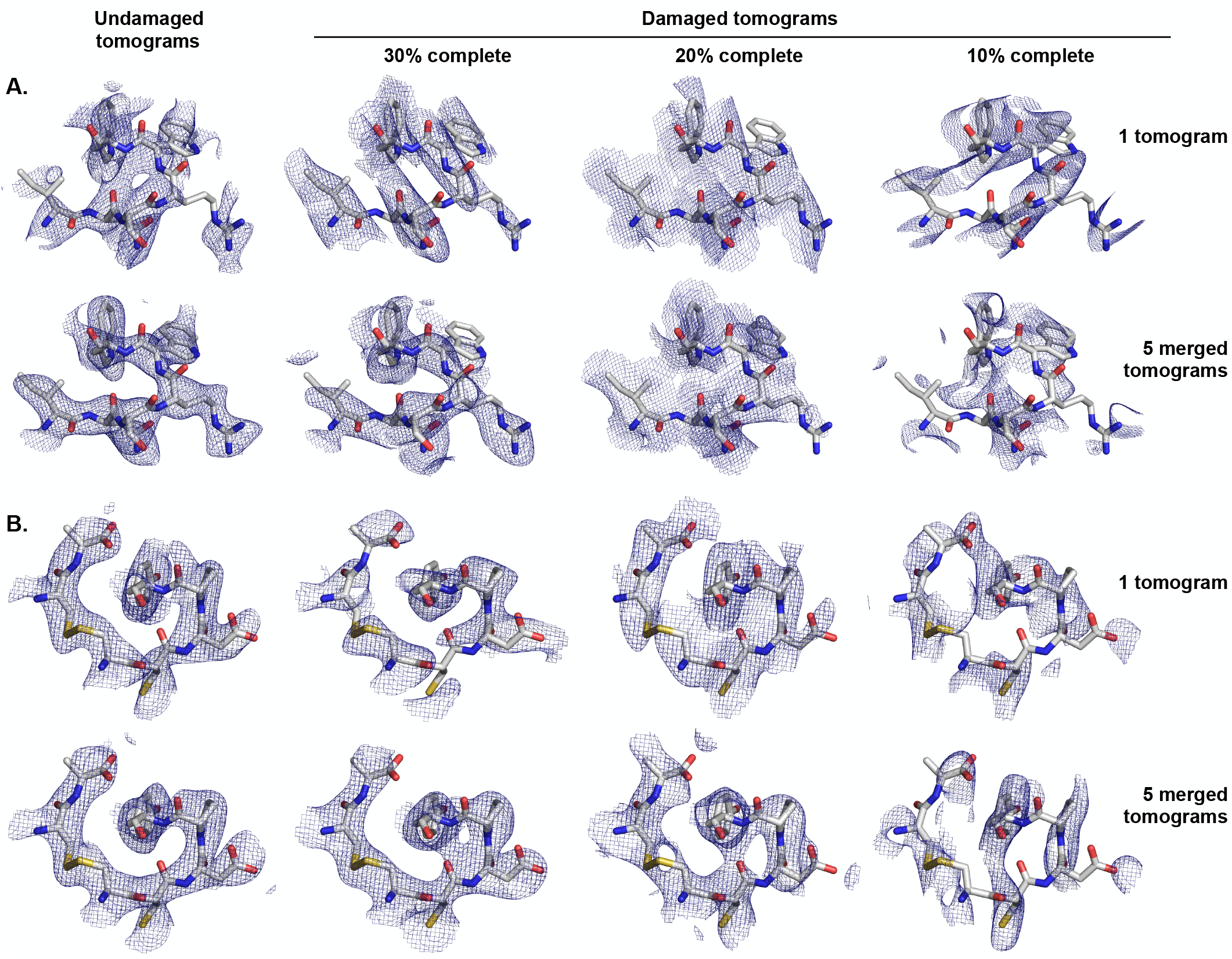
Merging tomograms with damage-induced loss of completeness. Density maps were computed from reflection data recovered from one representative undamaged tomogram or damaged tomogram with the specified *P*1 completeness (upper), and after merging five tomograms of the indicated type (lower). A subset of residues is visualized for the (A) triclinic and (B) orthorhombic crystal systems.

Although this analysis suggests that >10% completeness per dataset is required for recovery of the correct structure, loss of overall completeness is not the only hallmark of radiation damage. The progressive accumulation of damage alters the spatial distribution of the reflection data by reducing the number of reflections recorded late in the tilt-series, or at high tilt angles for a dose-symmetric scheme. In addition, the simulated damage introduced phase errors, though the loss of phase accuracy varied for the same amount of simulated dose (Table 1). To disentangle the impact of incompleteness, the angular spread of reflections across the tilt-range, and phase error on the ability to merge tomograms, we simulated radiation damage in reciprocal space according to the following model. For each simulated dataset, the crystal lattice was subjected to a random rotation in 3D. Structure factors were then computed to 3.3 Å, and the positions of Bragg peaks were predicted as a function of tilt angle; reflections in the missing wedge region were excluded. Reflection intensities were modeled as decaying with a B-factor that increased linearly with dose [62]:

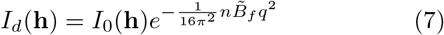

where *I*_0_ is the intensity of the undamaged reflection, *q* is the magnitude of the reciprocal space vector associated with reflection **h**, 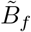 is a relative B-factor, and *n* is the image number in the tilt-series, with n=1,2,3… corresponding to tilt angles 0,−3,+3,−6… or 0,−2,+2,−4… for a dose-symmetric scheme spanning ±60 or ±40, respectively. The highestintensity reflections in the simulated tilt-range were retained to achieve the specified initial completeness. The phase origin was subjected to a random fractional shift (see Eq. 3), and phase errors were drawn from a normal distribution with the standard deviation chosen to achieve the target mean phase error. We assumed that the phase errors were dose-independent, since to our knowledge there are no models for how phase accuracy decays as a function of dose. This error model also assumes that the effects of tilt and the contrast transfer function, which modulate reflection phases with predictable behavior, can be corrected for [26]. The advantage of this damage model is that it allowed us to independently tune the phase accuracy, overall completeness, and angular spread of the reflection data across the tilt-range. Data were generated in this manner for the triclinic and orthorhombic systems used above in addition to a tetragonal crystal of proteinase K (PDB ID: 2ID8), which has a ~20-fold larger unit cell by volume and higher symmetry. Reflection data were then merged as described in Section IID.

Structure solution from tomograms of nanocrystals depends on finding the origin shift that correctly aligns the phases of multiple datasets. We assessed the accuracy of this merging procedure based on the merge error, the magnitude of the vector difference between the fractional origin shift estimated by our merging algorithm and the true shift required to align two datasets each subjected to a random phase shift. We considered the correct origin to be found when the merge error was less than twice the sampling interval used by the merging algorithm, corresponding to a fractional merge error of <0.03 for each crystal system. We then examined the dependence of this merge error separately on the phase accuracy, overall completeness, and the spatial distribution of the reflections of the datasets being merged. For the latter, we measured how unevenly the reflection data were distributed across the tilt-range using the Jensen-Shannon (JS) distance. This statistical metric measures the difference between a probability distribution of interest, *p*, and a reference probability distribution, *q*:

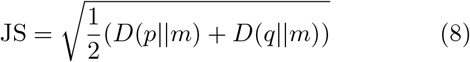

where *p* is the normalized distribution of tilt angles for the observed reflections, *q* is the normalized uniform distribution spanning ±60°, and 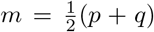. *D* is the Kullback-Leibler divergence given by:

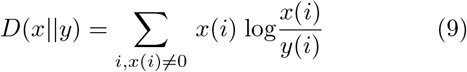

where the width of the angular bin *i* was set to 1°. Using our chosen reference distribution, the Jensen-Shannon distance measures how unevenly reflections are distributed across a full tilt-range of ±60°, and the score is independent of the dataset’s completeness. Datasets that span a narrower tilt-range than ±60° are penalized even when the reflection data are uniformly distributed since the normalized counts differ from the expected frequencies (Fig. S9). Radiation damage also increases the Jensen-Shannon distance due to loss of Bragg peaks at higher tilt angles as data collection progresses (Fig. S9).

The results from the three simulated crystal systems were pooled to examine trends that held across different space group symmetries and a range of unit cell volumes. We found that the initial phase error and the reflections’ angular spread across the tilt-series, as measured by the Jensen-Shannon distance, better predicted the likelihood of merging success compared to overall completeness (Fig. 6A). Provided that the reflections from individual tomograms were approximately evenly spread across the tilt-range (JS<0.18), the correct origin shift could be determined even in the presence of an average phase error of up to 40°. By contrast, even relatively high completeness (>50%, without accounting for space group symmetry) did not guarantee finding the correct phase origin in the presence of moderate phase errors (Fig. 6B-C). These trends argue for distributing the dose across as wide a tilt-range as possible to maximize the angular spread of reflections available from each tomogram. Although the real-space simulations indicated that a data collection scheme spanning ±40° with 2° increments would reduce phase errors relative to ±60° with 3° increments due to the narrower tilt increment (Table 1), these results predict that the modest increase in phase accuracy will be outweighed by the increased difficulty in correctly merging datasets due to the narrower angular distribution of the reflection data across the tilt-range.

**FIG. 6.**
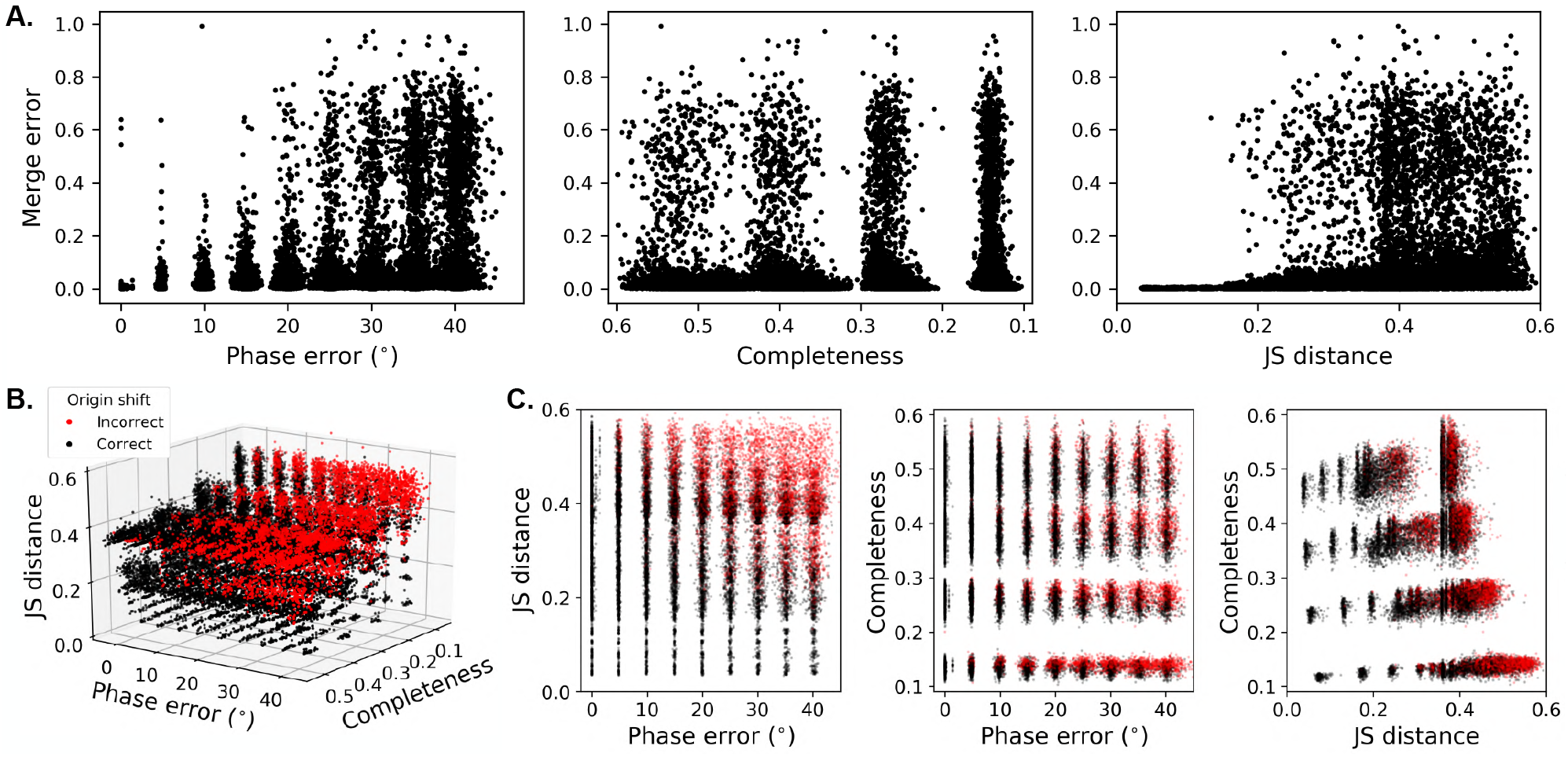
The dependence of merging success on phase accuracy, completeness, and the angular spread of reflection data. A total of 25,920 simulated datasets were generated across three crystal systems, spanning a range of *P*1 completeness, phase errors, and relative B-factors. Pairs of datasets with similar starting characteristics and in random orientations were merged. (A) The fractional merge error was computed as the magnitude of the vector error between the true and estimated fractional shifts required to position datasets on a common phase origin. This metric is shown as a function of the mean phase error (left), *P*1 completeness (center), and how unevenly spread the reflections were across the tilt-range (right) for the datasets being merged. The latter was estimated as the Jensen-Shannon (JS) distance to a uniform angular distribution spanning ±60° (see Eq. 8). Though 2880 pairs of datasets with a phase error of 0° were merged, the results are visually superimposed in the leftmost panel. (B) Merge errors are shown as a function of these three parameters or (C) the two indicated parameters of the datasets being merged. For (B) and (C), red and black respectively indicate datasets for which the incorrect and correct origin shift was determined by the merging algorithm. The error threshold was set to be twice the sampling interval used by the algorithm, such that the correct phase origin was found only when the fractional merge error was <0.03.

## IV. Discussion

Cryo-ET of protein nanocrystals could deliver a method of high-resolution structure determination that both retains experimental phases and circumvents the need for large crystals in conventional X-ray crystallography and high molecular weight proteins in SPR. Here we describe a data processing pipeline to solve structures from tomograms of nanocrystals. This workflow both leverages software from X-ray crystallography and provides new algorithms to handle challenges unique to tomographic data of crystalline specimens. These challenges are related to correctly extracting the reflection phases, including eliminating phase splitting due to imperfect periodicity, excluding partial reflections that result in phase errors of 180°, and extending the phase origin search procedure from 2D plane groups to 3D space groups. We validated this data processing scheme using simulated crystals and found that the recovered reflection information was accurate, yielding maps with a correlation coefficient to reference of ~0.9 after merging five tomograms without any structure refinement.

We also assessed the robustness of this pipeline to radiation damage, which limits the effectiveness of structure determination methods. Merging tomograms of crystals in different orientations is required to compensate loss of completeness, which results from both radiation damage and the missing wedge. We found that this merging procedure was especially sensitive to the angular spread of the reflection data across the tilt-range and phase errors but less so to the completeness of the individual tomograms. These simulations indicate a trade-off between a wider tilt-range to facilitate merging datasets and a finer tilt increment to reduce phase errors. Since including more datasets can overcome phase errors but not incorrect origin shifts, these results recommend data collection strategies that maintain a wide tilt-range rather than decreasing the sampling interval. We also predict that a wider tilt-range would be favored over finer sampling for high-resolution subtomogram averaging to similarly avoid aligning the missing wedge experienced by individual particles during alignment.

While this workflow addresses several data analysis issues specific to tomograms of nanocrystals, additional challenges remain. One critical issue is the optimal specimen thickness: thicker crystals increase signal-to-noise, but the accuracy of the signal suffers due to a larger defocus gradient and increased multiple scattering [26, 37]. We anticipate that the defocus gradient can be largely accounted for by a 3D correction of the contrast transfer function [63, 64], and microED has shown empirically that multiple scattering introduces only marginal (~5%) errors to the measured intensities for crystals 0.5-1 *μ*m thick [65, 66]. However, the impact of these effects on phase accuracy has not been characterized. Another common tomography challenge is the poorer image quality at high tilt-angles. Given the sensitivity of the merging procedure to the reflections’ distribution in reciprocal space, simulations that additionally account for loss of image quality at high tilt-angles may be useful for further refining data collection strategies from nanocrystals.

Despite these technical challenges, recent successes in cryo-ET of noncrystalline specimens support applying this technique to solve structures from nanocrystals to high resolution. In particular, subtomogram averages of purified immature HIV-1 Gag particles to 3.1 Å and *in situ* ribosomes to 3.7 Å demonstrate that high-resolution signal is available in tilt-series data [20, 21]. As with SPR, only particles with high molecular weight can be aligned with sufficient accuracy for this high-resolution information to be retained during subtomogram averaging. Alignment requires both determining the relative orientations of particles and positioning them on a common phase origin.

In comparison to SPR, the relative orientations of constituent proteins in a nanocrystal can be readily determined by indexing the coherent signal from the entire crystal. Even tiny nanocrystals will have a higher molecular weight than an individual HIV-1 Gag hexamer or ribosome, thereby extending the reach of cryo-ET to smaller proteins. In comparison to X-ray crystallography, both microED and cryo-ET of nanocrystals require a wider angular range of reflection data for indexing due to the negligible curvature of the Ewald sphere using electron radiation. Though successful indexing of individual electron diffraction stills has been reported [67], in general a tilt-range of ±20° is considered necessary to index microED data [68]. Cryo-ET data pose additional challenges, as imaging but not diffraction data are negatively impacted by several factors including the defocus gradient, translational drift, and loss of coherency due to inelastic scattering. These effects have been predicted to cumulatively reduce the signal-to-noise ratio of projection images by over 90% compared to diffraction stills collected from the same crystal, with increasing loss at higher resolution and for thicker crystals [69]. The diminished signal-to-noise will result in fewer observed reflections when collecting data in imaging versus diffraction mode for a fixed dose or exacerbate radiation damage if the dose is increased to compensate. Either scenario could make indexing more difficult for cryo-ET compared to microED data. Here we adapted algorithms from the crystallography software package DIALS to index the Fourier transforms of simulated nanocrystal tomograms [41, 42]. We found that indexing was robust even for simulated tomograms with low completeness.

While the use of nanocrystals facilitates determining relative particle orientations, the low molecular weight of the constituent proteins makes finding a common phase origin – the other step of particle alignment – more challenging. High-resolution subtomogram averaging has exclusively targeted high molecular weight proteins, and with two exceptions, isolated particles in solvent [19–21, 70]. These characteristics facilitate finding a common phase origin from each particle’s center of mass. For nanocrystals composed of small and densely-packed proteins, a more extensive search over the unit cell volume is required to position datasets on a consistent phase origin. Finding a common phase origin to merge datasets is critical for improving the accuracy of the recovered reflection data and overcoming low completeness.

In the future, we anticipate that cryo-ET of nanocrystals could enable structure determination from disordered crystals, which are typically discarded during diffraction experiments. In crystals, disorder attenuates the high-resolution signal and has been observed at length scales relevant for nanocrystals [25, 35]. In contrast to diffraction methods, imaging permits spatially characterizing and computationally correcting such disorder. One approach is to denoise the tomogram in Fourier space, followed by computationally “unbending” lattice distortions using cross-correlation analysis between the denoised and original tomograms. Lattice unbending overcame the resolution-limiting effects of disorder in 2D electron crystallography, in which specimens were frequently bent or wrinkled [26, 27]. Alternatively, subtomogram averaging could be performed in real space [38]. This technique has achieved subnanometer reconstructions of pleomorphic viruses with imperfect helical symmetry and could prove similarly useful for solving structures from tomograms of disordered nanocrystals [71]. In addition to extending the high-resolution limit of structure determination, the ability to spatially characterize disorder could provide insights into both the organization of proteins that form ordered arrays *in vivo* and defects in these biological crystals [72–74].

In addition, we anticipate that the development of a hybrid method involving microED and cryo-ET of nanocrystals could become routine. This approach would combine diffraction intensities with low-resolution phases from experimental images to solve an initial structure, followed by phase extension to the high-resolution limit of the microED data [4, 75]. Combining these techniques will be facilitated by the fact that sample preparation is shared between the techniques, and data can be collected on the same instrument [10]. Implementation of both disorder corrections and a hybrid approach with microED will be critical to realize the full potential of cryo-ET of nanocrystals for high-resolution structure determination.

## Code availability

The code developed to process tomograms of nanocrystals is available at https://github.com/apeck12/cryoetX.

## Acknowledgments

We thank Lauren Ann Metskas and Florian Schur for valuable discussions, in addition to David Stokes and Steven Ludtke for advice on the phase splitting phenomenon. A.P. is The Mark Foundation for Cancer Research Fellow of the Damon Runyon Cancer Research Foundation (DRG 2361-19). This work was supported by NIH grants R35 GM122588 (to G.J.J), AI150464 (to G.J.J), and GM117126 (to N.K.S).

