## Supplemental Figures for "Challenges in solving structures from radiation-damaged tomograms of protein nanocrystals assessed by simulation"

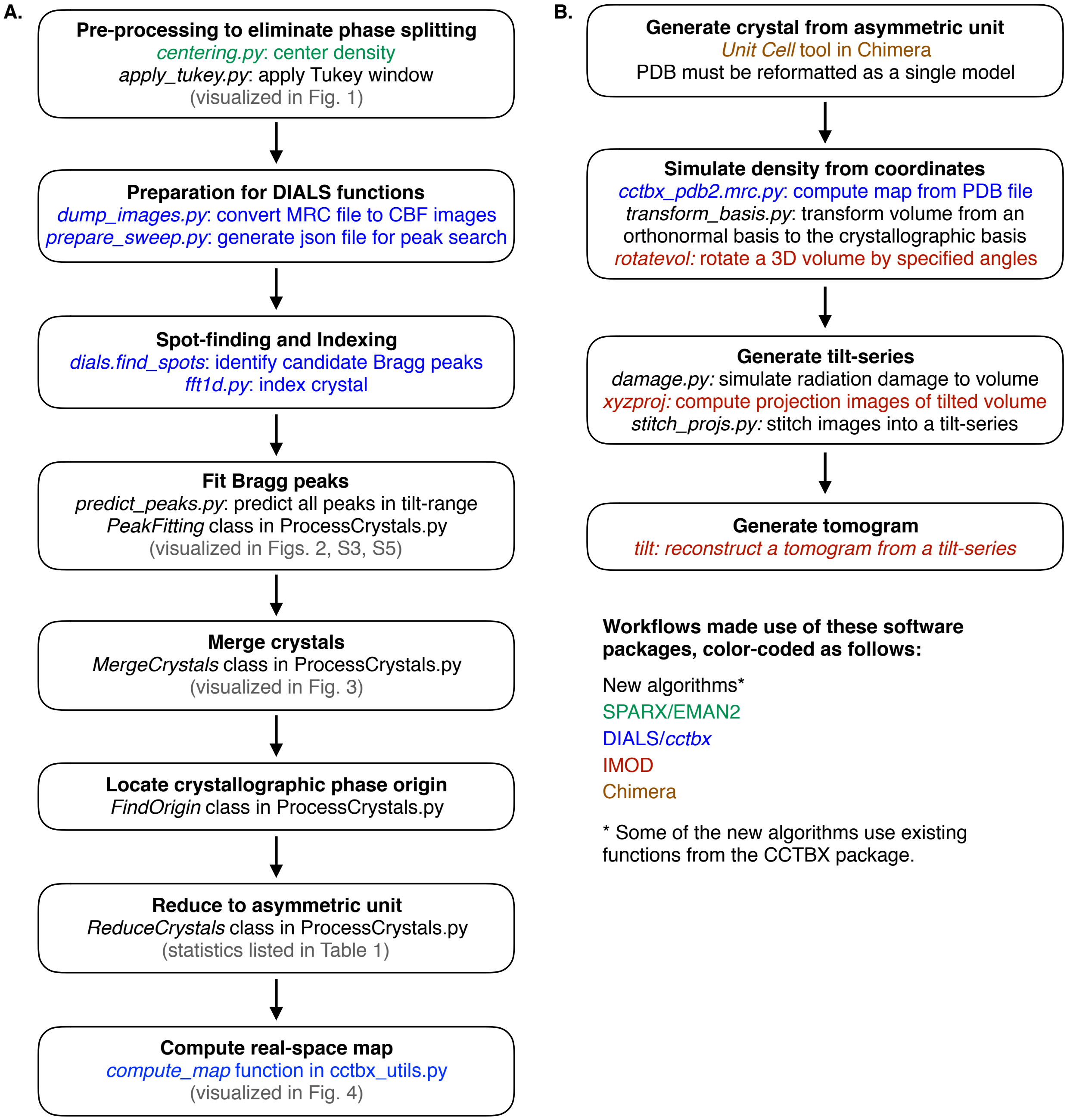


FIG. S1. **Flowcharts of data processing pipeline and data simulation.** The workflows used to (A) process and (B) simulate tomograms of protein nanocrystals are outlined. Steps that relied heavily on functions from the SPARX or EMAN2, *cctbx* or DIALS, IMOD, and Chimera software packages are respectively shown in green, blue, red, and brown. New algorithms that use existing functions from the *cctbx* library but are specific for tomographic data of nanocrystals are in black. All Python scripts are available at: https://github.com/apeck12/cryoetX.

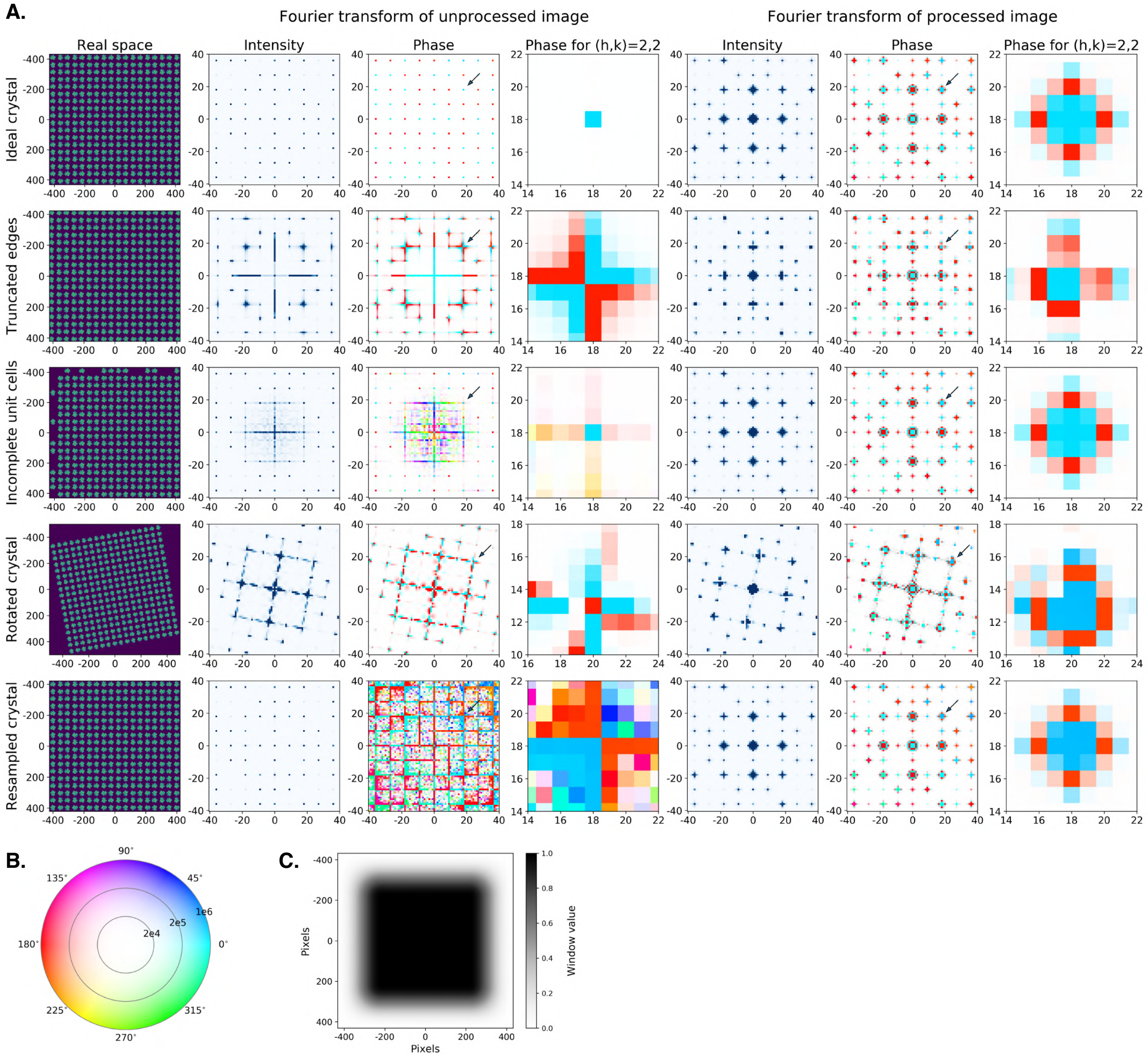


FIG. S2. **Imperfections result in phase splitting at Bragg peak positions.** (A) All crystals were generated from a 2D unit cell consisting of 4 blobs related by C4-symmetry. The leftmost column shows the real space density, while the second and third columns respectively show a central region of the intensities and phases from each crystal's Fourier transform. The black arrow indicates a representative Bragg peak, $h,k$=(2,2). The phases in the vicinity of this peak are shown in inset in the fourth column. A tapered cosine (Tukey) window function was applied to the real space crystal density, which was then centered within the image boundaries before being shifted to the top left corner and computing the Fourier transform. Columns 5-7 respectively show the intensities, phases, and the phases specifically in the vicinity of the (2,2) peak after these image processing steps. In all phase plots, pixel color corresponds to the phase value while saturation is determined by the intensity value. For the *ideal crystal* (row 1), the unit cell was tiled exactly along the x and y coordinate axes to generate a finite but otherwise ideal lattice. For the crystal with *truncated edges* (row 2), a region spanning ten pixels wide along each border of the ideal crystal was removed. For the crystal with *incomplete unit cells* (row 3), asymmetric units that border two of the ideal crystal's edges are randomly retained or omitted. For the *rotated crystal* (row 4), the ideal crystal is rotated counterclockwise by 10°. For the *resampled crystal* (row 5), the ideal crystal is interpolated onto a grid in which the unit cell does not span an integer number of pixels. (B) A color wheel shows how pixels are visualized in columns 3-4 and 6-7 of (A) and Figs. 1-2 of the main text. Color and saturation respectively indicate the phase and intensity values of each pixel. (C) The tapered cosine window applied to the real space crystal is shown. A 3D version of this window function is used for the simulated protein crystals presented in the main text.

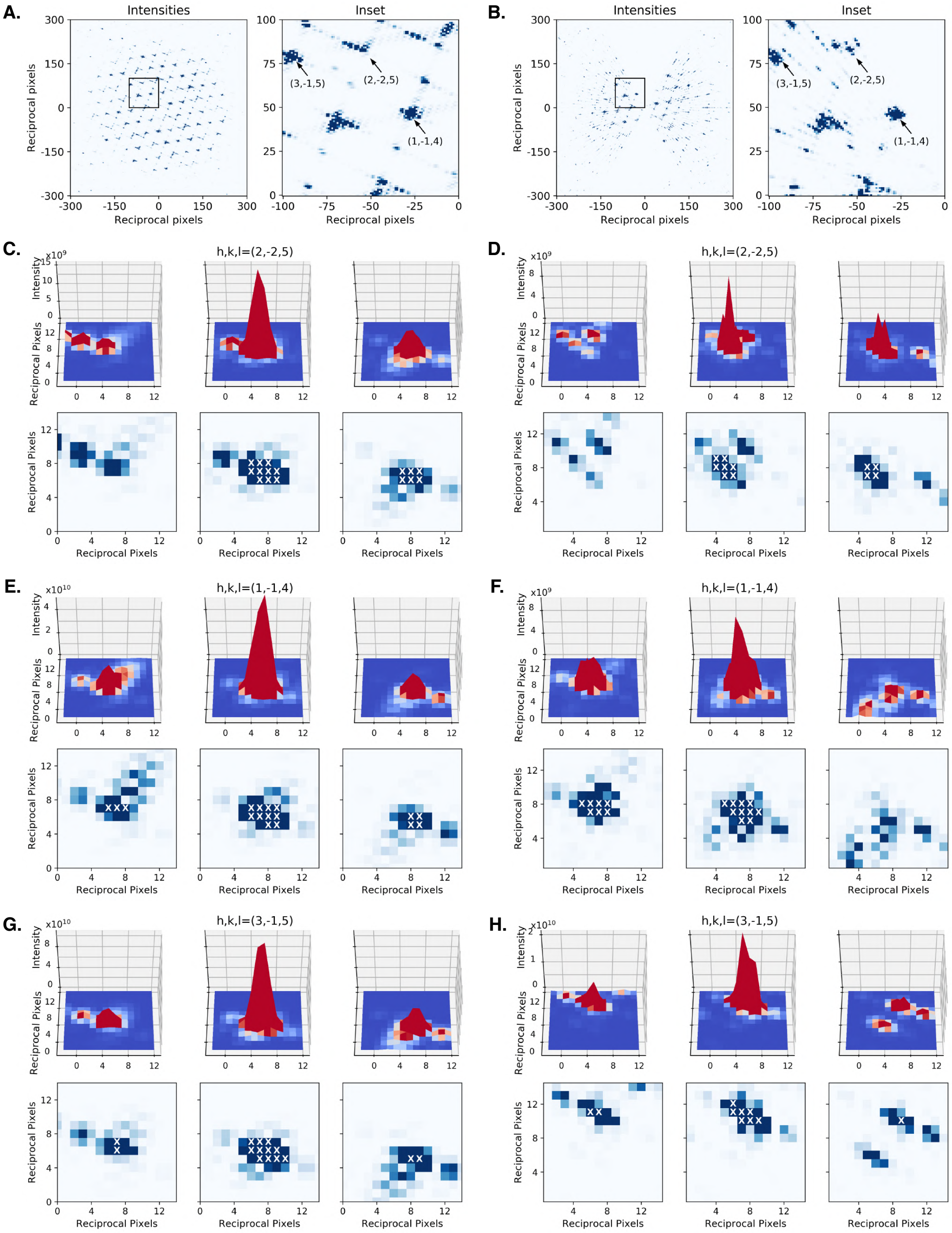


FIG. S3. **Anistropic background due to tomographic sampling.** Intensities from a slice through the Fourier transform are shown from (A) a simulated crystal and (B) a tomogram generated from this crystal with 3° increments between tilts. The boxed region is shown in each inset. In (C-H), the indicated reflection is visualized, with the middle panel showing a slice through the reflection's center. The panels on the left and right show slices one reciprocal pixel above and below this plane, respectively. In each, the reciprocal pixels assigned to the Bragg peak are marked with an X in the lower panel. These reflections are shown from the intact crystal in (C,E,G) and the tomogram in (D,F,H). Each reciprocal pixel spans 1 $Å^{-1}$, and the resolution of the (2,-2,5), (1,-1,4), and (3,-1,5) reflections is respectively 5.8, 9.2, and 4.7 Å.

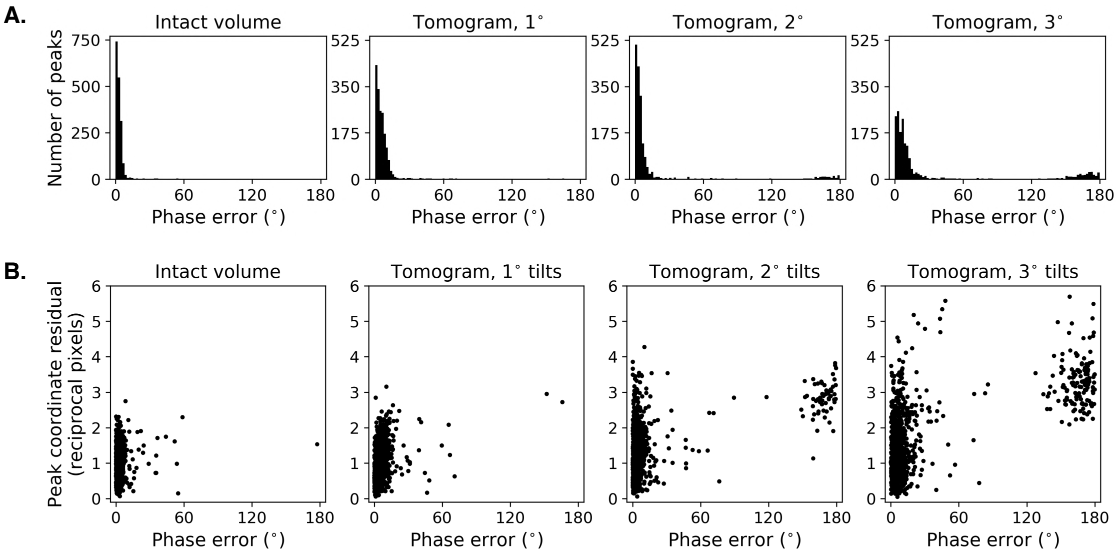


FIG. S4. **Phase accuracy decreases as the tilt increment between projection images increases.** A simulated crystal (intact volume) was projected into tilt-series spanning a $\pm 60^\circ$ tilt-range using 1, 2, or $3^\circ$ increments between images. Tomograms were generated from the tilt-series, and the Bragg peak phases were extracted from the tomogram's Fourier transform and shifted to a crystallographic phase origin. (A) The distributions of residuals between estimated phases from the indicated dataset and reference phases computed from the atomic model are shown. (B) The distance between the observed and predicted coordinates of each Bragg peak center is plotted as a function of the peak's phase error for the indicated dataset.

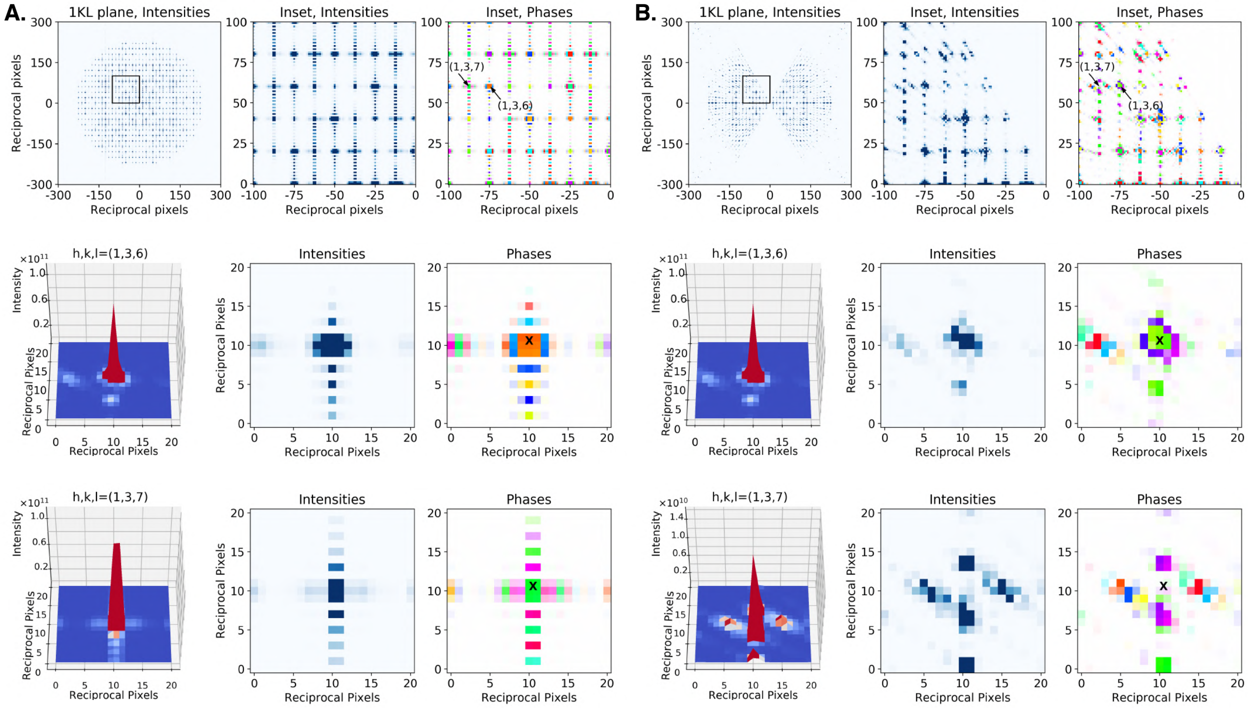


FIG. S5. **Representative Bragg peaks show that a 3° tilt increment results in poor sampling of peak centers.** *Upper*: Intensities from the 1KL plane are shown in the left panels from the Fourier transforms of (A) an intact volume and (B) a tomogram of a simulated crystal. The center and right panels respectively show the intensities and phases from the subregion indicated by the black box. *Middle*: The (1,3,6) reflection is visualized as a representative peak whose center is well-sampled in both cases, with a difference of <1 reciprocal pixel between the ideal Bragg peak position (marked by an X in the phase plot) and the center estimated by peak-fitting. After shifting each simulated crystal to the reference phase origin, the phase error for this reflection was 0.4° and 7° for the (A) intact volume and (B) tomogram, respectively. *Lower*: The (1,3,7) reflection is visualized as a representative peak whose center is poorly sampled in the tomogram due to the 3° increment between tilt images. In the case of the (A) intact volume, the ideal and estimated peak positions coincide, with a discrepancy of <1 reciprocal pixel, and the phase residual to reference is 0.4° after the crystal is shifted to the reference origin. In the case of the (B) tomogram, there is a coordinate error of 3 reciprocal pixels between the ideal (indicated by an X) and observed peak positions, resulting in a phase residual to reference of 173° for this reflection. Each reciprocal pixel spans 1 $Å^{-1}$, and the resolution of the (1,3,6) and (1,3,7) reflections is 5.9 and 5.4 Å, respectively.

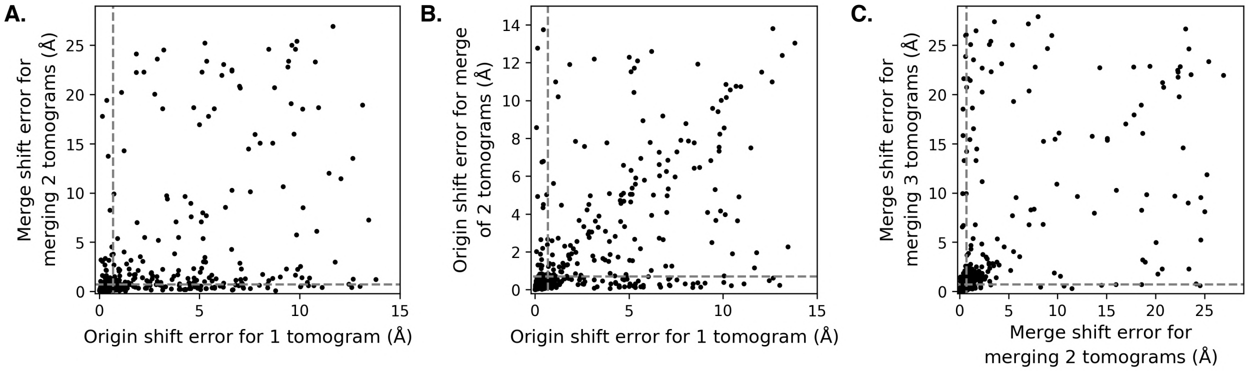


FIG. S6. **Comparison of the robustness of merging datasets before locating a crystallographic origin and vice versa.** Simulated datasets for a $P2_12_12_1$ model crystal were generated with a range of initial completeness (10-40% prior to accounting for symmetry) and mean phase errors (0-40°). (A) Each point compares the error in determining the crystallographic phase origin (origin shift error) for a single dataset and the error in merging this dataset and a second dataset (merge shift error). All datasets were positioned on a random phase origin. There were 4% more cases of successfully merging two datasets than finding the correct crystallographic origin for a single dataset. In (B), the error in determining the crystallographic phase origin is compared for a single tomogram and the merge of two tomograms. There were 8% more successful cases for the latter. In (C), the error for merging two and three tomograms is compared. The number of successful cases in each case was similar. In each plot, the dashed lines indicate the threshold below which the correct merge or origin shift was found, and the threshold is based on the sampling step used during the search procedure. The maximum possible error is lower for the origin shift because there are four valid crystallographic origins for $P2_12_12_1$ crystals, but only one correct origin for merging two datasets that are not necessarily positioned on a crystallographic origin.

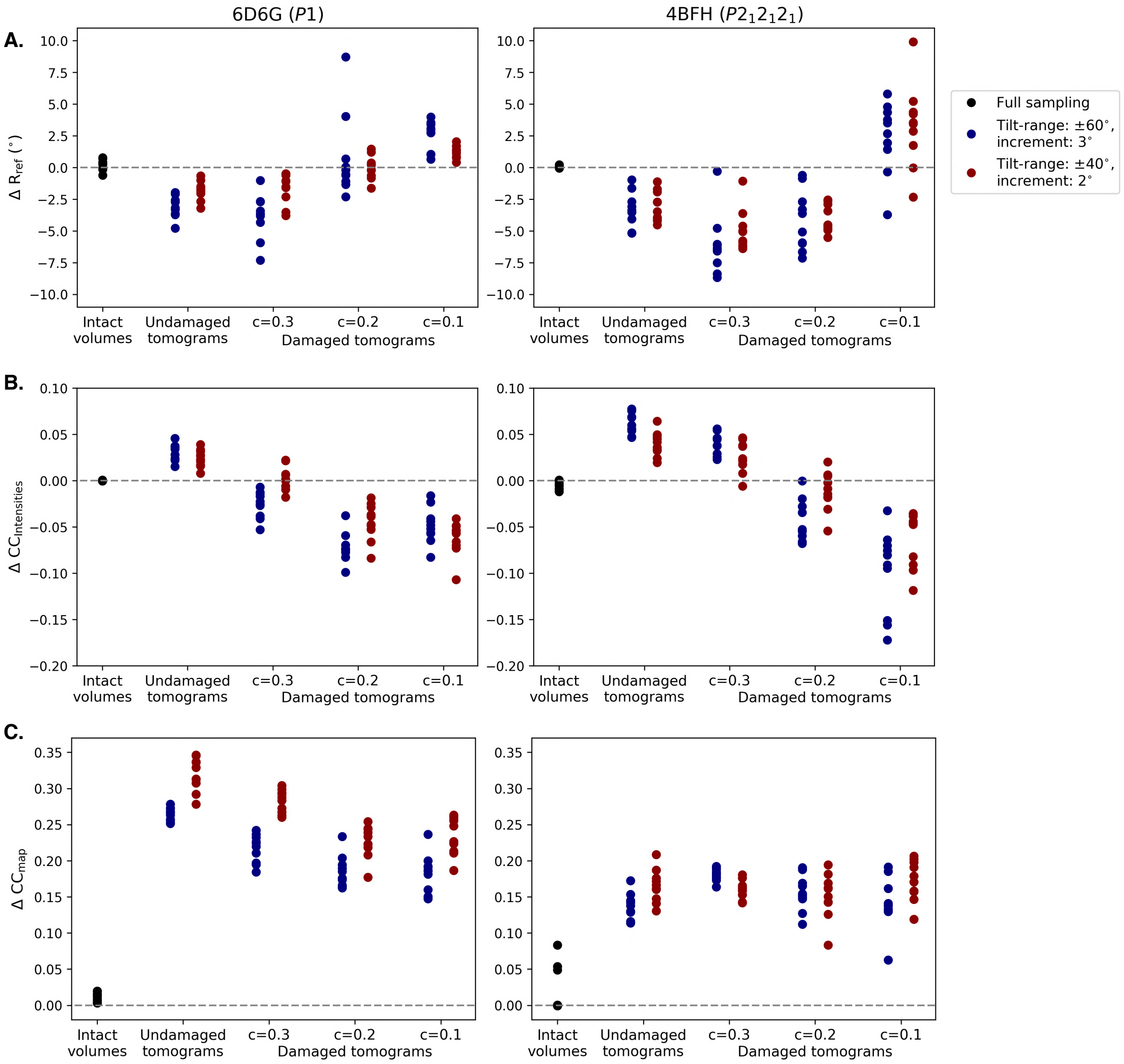


FIG. S7. **Changes in data quality from merging multiple tomograms.** The changes in the (A) mean phase error relative to the reference phases, (B) cross-correlation between the logs of the estimated and reference intensities, and (C) cross-correlation with the reference map from merging the indicated type of data are shown. Each point represents the difference between the merged data and the average value of the five tomograms included in the merge. Results are shown for the simulated triclinic (left) and orthorhombic (right) crystal systems, and compared for intact volumes (black) and tomograms generated from either a $\pm 60^\circ$ tilt-increment (blue) or a $\pm 40^\circ$ tilt-increment (red). For the damaged tomograms, "c" denotes the fractional $P1$ completeness of the individual tomograms; for comparison, undamaged tomograms were characterized by a $P1$ completeness of 0.4. For (A), points below the dashed line indicate datasets for which merging improved the phase accuracy, while for (B) and (C), points above the dashed line indicate that merging improved the accuracy of the reflection intensities and real-space maps.

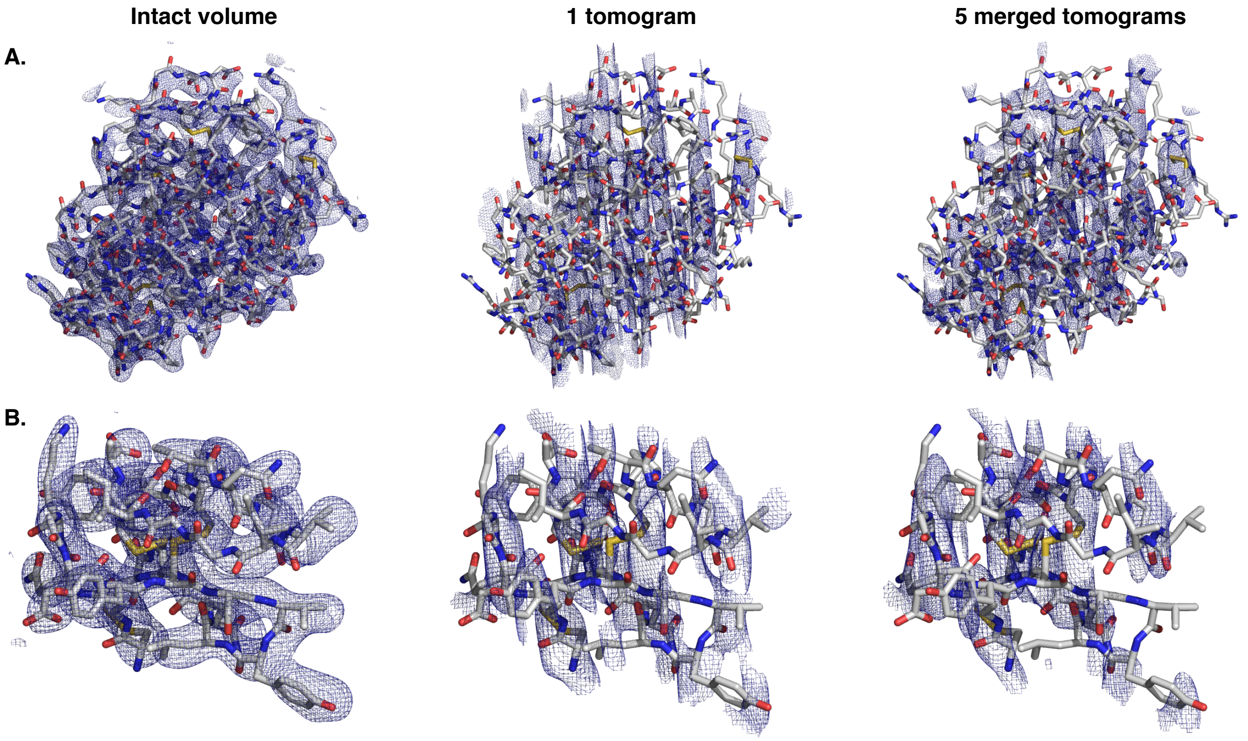


FIG. S8. **Merging tomograms with low completeness aligns the missing wedge.** Density maps computed from the reflection data extracted from an intact volume (left), a single tomogram with a $P1$ completeness of 10% (center), or the merge of five 10% complete tomograms (right) are shown for the (A) triclinic and (B) orthorhombic model systems.

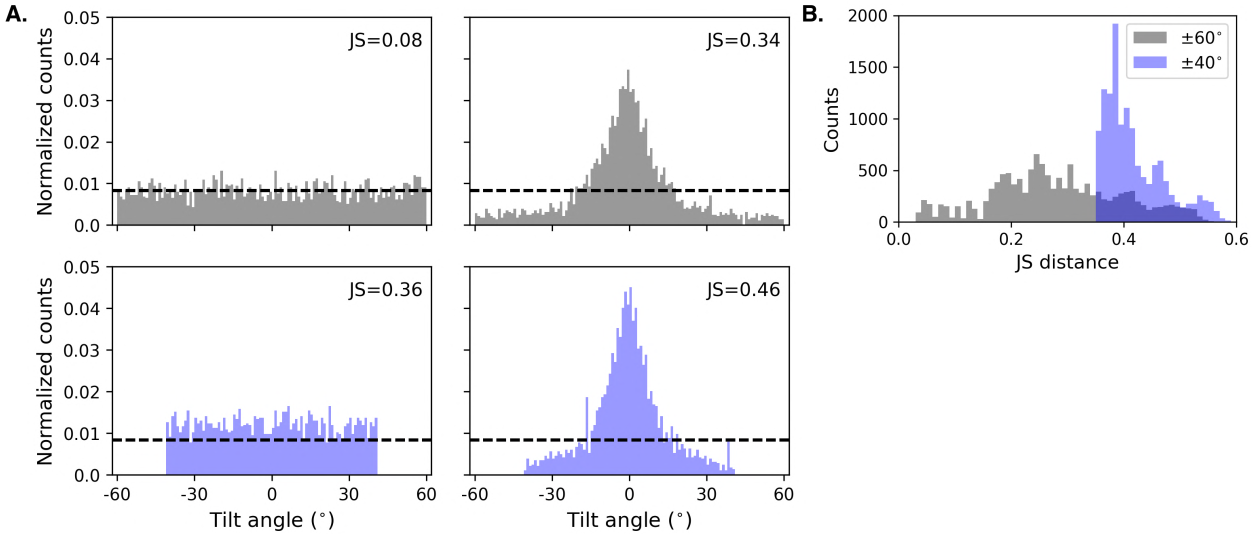


FIG. S9. **The Jensen-Shannon distance measures the angular spread of reflections over the tilt-range.** The Jensen-Shannon (JS) distance is a statistical metric that measures the difference between a distribution of interest and a reference probability distribution. Here a uniform angular distribution across ±60° was used as the reference distribution, such that the score measured how unevenly distributed in reciprocal space the observed reflections were across this possible tilt-range. (A) The angular distributions of reflections from representative datasets simulated using a ±60° (upper) or ±40° (lower) tilt-range and a relative B-factor of 0 Å (left) or 600 Å (right) are shown for a tetragonal crystal (PDB ID: 2ID8). For each dataset, the overall $P1$ completeness is 10%. The dashed line indicates the expected normalized counts for a uniform distribution spanning ±60° with an angular bin width of 1°. The Jensen-Shannon score for each dataset is noted in the upper right corner. (B) Merging was attempted for 12,960 pairs of simulated datasets that were generated across three crystal systems, spanning a range of phase errors, initial completeness, relative B-factors. The distributions of the Jensen-Shannon distances for simulated datasets generated with either a ±60° (grey) or ±40° (blue) tilt-range are compared.
